# Moving by thoughts alone? Amount of finger movement and pendulum length determine success in the Chevreul Pendulum Illusion

**DOI:** 10.1101/841445

**Authors:** Debora Cantergi, Bhuvanesh Awasthi, Jason Friedman

## Abstract

Hand-held pendulums can seemingly oscillate on their own, without perceived conscious control. This illusion, named after Chevreul, is likely a result of ideomotor movements. While this phenomenon was originally assumed to have a supernatural basis, it has been accepted for over 150 years that the movements are self-generated. However, until now, recordings of the small movements that create these oscillations has not been performed. In this study, we examined the mechanism that produces these significant involuntary oscillations using a motion capture system. We determined that the Chevreul pendulum illusion is produced when the fingers holding the pendulum generate an oscillating frequency close to the resonant frequency of the pendulum. At an appropriate frequency, very small driving movements of the arm are sufficient to produce relatively large pendulum motion. Further, subjects that tended to move their fingers more were more successful in producing the illusion.

## Introduction

Can humans cause movements in external objects unintentionally? Are there motor actions that result from thoughts or mental images and are potentially instantiated independently of conscious engagement? Once attributed to external spirits, non-conscious motions of the hand-held pendulum have been attributed to ideomotor phenomenon. Ideomotor theory posits that actions are represented by their perceivable effects (Shin, Proctor, & Capaldi, 2010). This theory has been used as a way of explaining how voluntary movements can occur when one is not consciously aware of making movements. A classic example of this is when a hand-held pendulum will start moving without the holder feeling like they are performing any movement.

In 1808, Antoine-Claude Gerboin from Strasbourg School of Medicine described his observation of how the hand-held pendulum would move mysteriously when the person held it over certain substances. In the 1830s, a French scientist, Michel Eugene Chevreul studied the movement of such a hand-held pendulum in several situations (Chevreul, 1854; Jastrow, 1962), verifying that the movement decreased when the arm was being supported at the hand in contrast to the arm being externally supported at the shoulder, and that the oscillations were sight dependent. Chevreul postulated that imperceptible muscle activations were responsible for the pendulum’s first oscillations, which increased (but were still imperceptible) under the influence of visual feedback (Borowczyk, 2011). Although nearly 200 years have passed since this and similar examples of ideomotor behavior, such as Ouija board spelling were explained (Spitz, 1997), people are still being deceived by modern versions of ideomotor behavior, including tragically in facilitated communication (Burgess et al., 1998).

There have been several attempts to better understand the Chevreul pendulum illusion. In the 1970s, Easton and Shor used photogrammetry (Easton & Shor, 1975, 1976) and confirmed that sight is an important factor in determining when the pendulum will oscillate (Easton and Shor, 1975), and that as more attention is directed toward the pendulum, the more it oscillates (Easton and Shor, 1976). They also confirmed that restraining the arm at the wrist decreased the pendulum oscillation and demonstrated that out of 75 subjects, only 60 were able to create the illusion (Easton and Shor, 1976). Hypnosis research uses Chevreul’s pendulum illusion as a tool for testing a patient’s response to the technique, with patients that are not able to move the pendulum generally being unresponsive to hypnosis (Karlin et al., 2007).

The studies of Easton and Shor, however, were not sensitive enough to describe how the subjects generate the pendulum motion. Despite the early research and ongoing public fascination with these phenomena, there is relative paucity of research examining the mechanistic accounts of ‘automatic’ pendulum oscillations.

A pendulum has a resonant frequency that is primarily dependent on its length. The maximum oscillation amplitude for a hand-held pendulum will be achieved if the driving frequency (i.e. the frequency of oscillations of the hand) is equal to the resonant frequency (Newburgh, 2004). Thus, in order to make the pendulum oscillate significantly, the subjects are required to oscillate the pendulum-holding fingers at a frequency close to the resonant frequency. Here, we set out to examine this relationship between resonant frequency and the possible ‘ability’ of the subjects to ‘will’ the movements in the pendulum. In this study, we chose pendulums that had natural frequencies of approximately 0.7, 1.05 and 2.0 Hz. This means that higher-frequency physiological tremor, e.g. 3-5 Hz for the elbow and 8-12 Hz for the finger (Hallett, 1998), should not produce significant pendulum motion, rather, it will require oscillations generated “voluntarily”. Using motion capture equipment, we aim to investigate what movement subjects make in order to produce the pendulum illusion, determine how (i.e. with which body parts) subjects produce oscillations, and to examine at which frequencies subjects are able to generate pendulum oscillations.

## Methods

### Subjects

Thirteen right-hand dominant subjects (9 females), with normal or corrected vision took part in the study. The sample size was based on a pilot experiment (Cantergi & Friedman, 2018), where we observed a 30% difference in arc length between conditions, and a standard deviation of approximately 50% of the mean, which gives a sample size of 13 for a one-sided signed-rank test (Faul, Erdfelder, Lang, & Buchner, 2007). The experimental protocol received ethical approval from the Tel Aviv University Institutional Research Board, and subjects signed an informed consent form before starting the experiment. Subjects received payment of 40 shekels (approximately $11) for participating in the experiment.

### Procedure

Subjects stood at a marked location, with their hand outstretched in front of them. A magnetic motion capture system (Polhemus Liberty), sampling at 240 Hz with 8 sensors was used to record the movement of the pendulum and the right upper limb, using the Repeated Measures software (Friedman, 2014). One sensor was used as the pendulum, the other sensors were placed on the thumb, index and middle fingernails, on the back of the palm, on the forearm, on the upper arm, and on the shoulder. Each recording was for 120 seconds. A short period of rest between attempts was provided. Before starting the recordings, subjects were instructed to perform upper arm movements (each lasting 5 secs) of one of the seven degree of freedom in each case, to allow calculation of the joint centers and joint angles.

The subjects first held their arm outstretched without the pendulum. In the second trial, they held the wire 40 cm from the sensor, with no attempt made to move it. In the third, fourth and fifth trials, they held the pendulum and attempted to cause it to move (without consciously moving their hand). This was performed with a 40 cm pendulum (as before), then with a pendulum half the length (20cm), and double the length (80cm). Finally, in the sixth trial they repeated the attempt to move the 40cm pendulum, but without visual feedback.

In order to determine the resonant frequency of the pendulum, the wire was attached to a stand, pulled to 45 degrees, and left to swing on its own. The resonant frequency was then calculated from the reciprocal of the mean distance between the extreme values on one side. The resonant frequencies for the 20cm, 40cm and 80cm pendulums were found to be 2.00 Hz, 1.05 Hz and 0.70 Hz respectively.

### Data Analysis

The position and orientation data from the 8 sensors were smoothed using a 4^th^ order, twoway low-pass Butterworth filter, with a cutoff of 5 Hz. Joint centers and subsequently the joint angles were calculated using a previously described technique (Biryukova, Roby-Brami, Frolov, & Mokhtari, 2000). A fast Fourier transform (FFT) was performed on each trial on the thumb velocities. As the pendulum is likely to move close to its resonant frequency due to any finger movements, we classified successful trials as those where both the pendulum moved a considerable amount (peak-to-peak amplitude greater than 10°) and the thumb FFT showed its largest peak close (defined as less than 15%) to the resonant frequency of the pendulum.

For each trial, we calculated the arc length, defined as the total amount of movement of the thumb, relative to the shoulder, i.e., we calculated the unsigned distance between each two time points, and then summed them. We also determined the relative contributions of the shoulder, elbow and wrist to left-right movements of the thumb in the relevant frequency range (within 0.2 Hz of the pendulum frequency). After calculating the joint angles at the shoulder, elbow and wrist, we considered only the component responsible for rotations about the vertical axis (i.e. those that cause left-right movement). We then computed the joint velocity (by taking the derivative), and computed the tangential velocity at the thumb (by multiplying the joint velocity by the joint-thumb distance). We computed the relative contribution by dividing the area under the relevant region of the FFT for one joint by the sum of the areas for the three joints.

We compared the arc lengths between successful and unsuccessful participants using a onesided Wilcoxon signed rank test, under the hypothesis that successful participants will move more than unsuccessful participants. We presented the W statistic for the test, which is the sum of the ranks of positive differences, along with the approximate 95% confidence intervals (Altman, Machin, Bryant, & Gardner, 2000). Statistical analyses were performed using Matlab 2018b (Mathworks).

## Results

Some of the subjects were successful in achieving the pendulum illusion. The success was greatest using the 40 cm pendulum with vision (62%), lower success rates were achieved with the longer (46%) and shorter pendulums (8%), and when no visual feedback was allowed (31%). Four subjects (31%) were unsuccessful in all tasks. An example of a successful and unsuccessful subject for the 40cm pendulum task with vision is shown in Figure 1.

**Figure 1.**
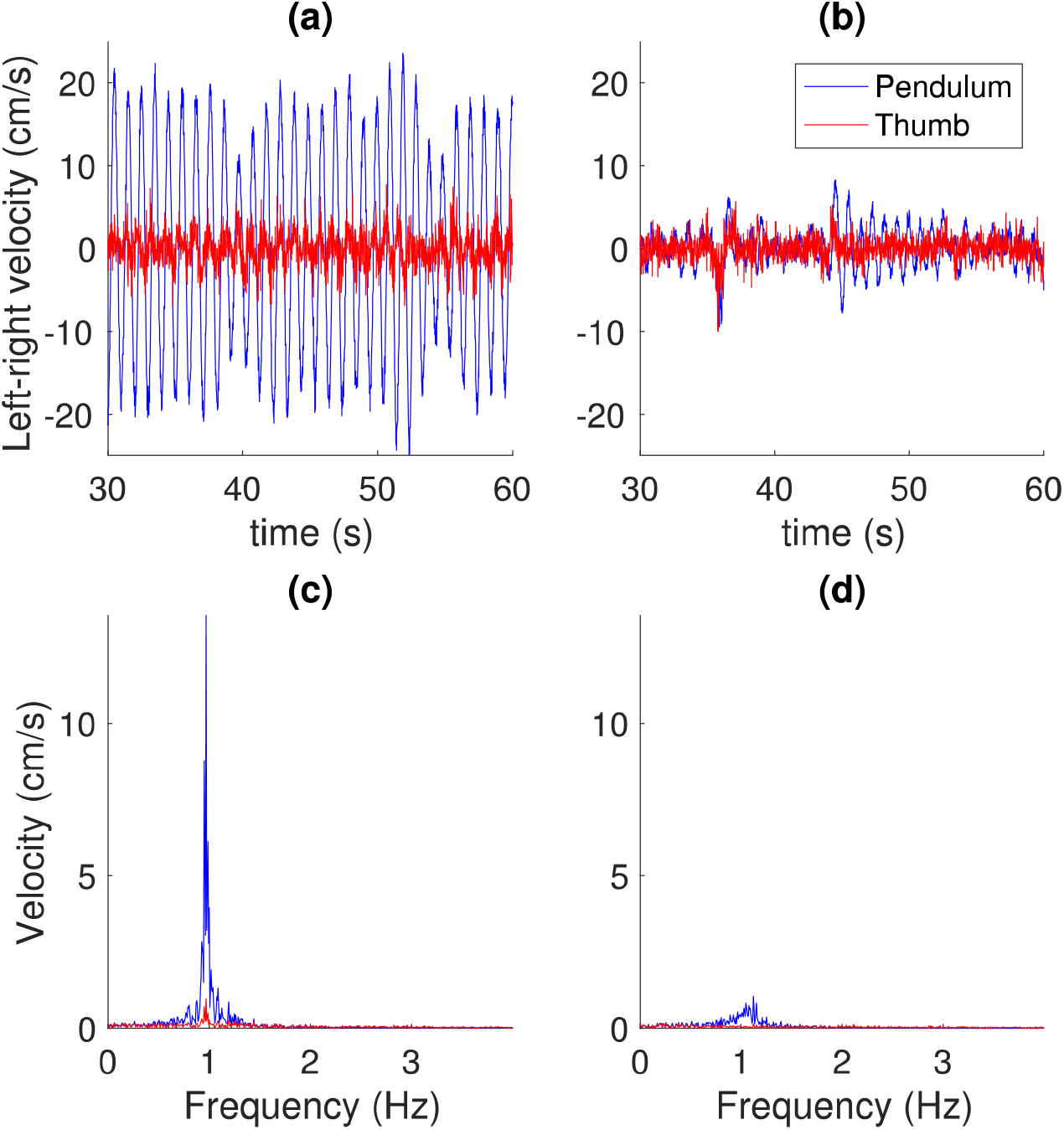
Example of a successful (left column) and unsuccessful (right column) subject. (a) and (b) show the left-right velocity of the pendulum and the thumb (for a 30 second portion of the trial). (c) and (d) show the Fourier transforms (using data from the whole 2 minutes) – for the successful subject (c), a clear aligned peak can be observed at approximately the resonant frequency of the pendulum for both the pendulum and the thumb, whereas no peak is present for the thumb or pendulum at this frequency for the unsuccessful subject (d).

We computed the arc length of the thumb in the different conditions, shown in Figure 2. When attempting to move the pendulum at 40cm, those successful in moving the pendulum showed greater arc lengths (median 249 cm, CI = [140, 881] cm) than those who were unsuccessful for that condition (median 169 cm, CI = [122, 180] cm; Wilcoxon rank sum test: W = 19, p=0.001), as were those at 80 cm (successful: median 273 cm, CI = [152, 779] cm; unsuccessful: median 159 cm, CI = [89, 283] cm; W=34, p=0.017) and in the condition with no vision (successful: median 253 cm, CI = [203, 301] cm); unsuccessful: median 177 cm, CI = [171, 224] cm; W=50, p=0.025). Further, success in moving the 40 cm pendulum was shown by subjects who had a greater arc length in the first trial, when they held their arm out without holding the pendulum (successful: 258 cm, CI = [114, 739] cm; unsuccessful: median 157 cm, CI = [137, 218] cm, W=23, p=0.046).

**Figure 2.**
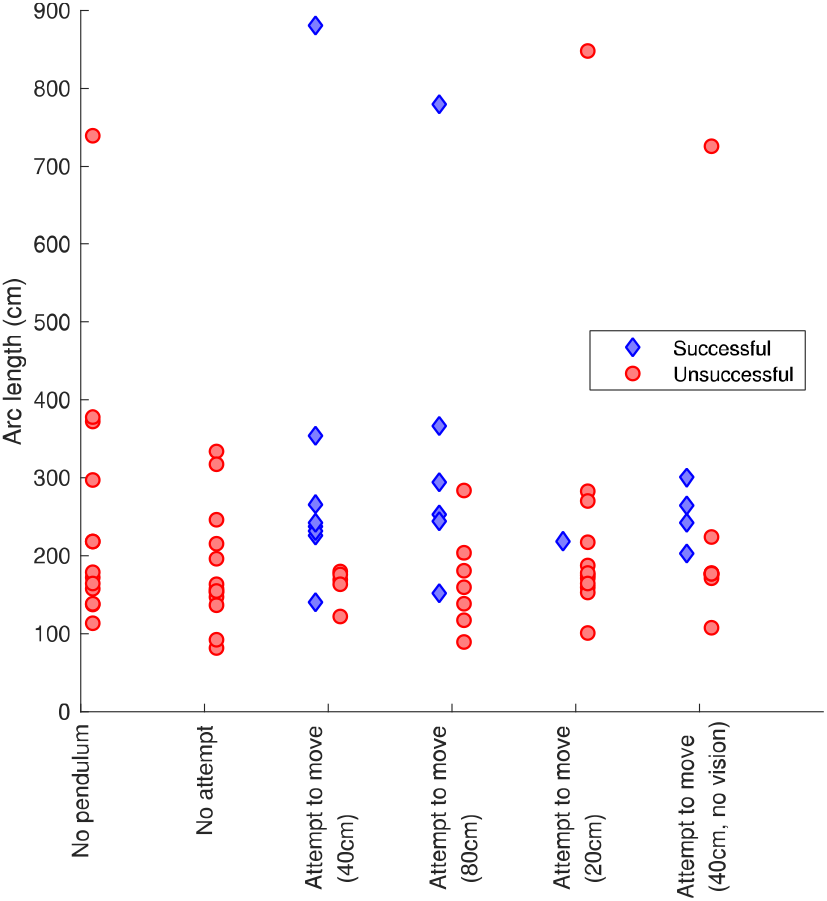
Comparison of arc lengths (over two minutes) in the six different conditions. Blue dots indicate successful trials, red dots unsuccessful trials

We also examined how subjects produced the illusion, i.e. the effect of shoulder, elbow and wrist velocities on left-right thumb velocity in the relevant (± 0.2 Hz from the pendulum frequency). The relative contributions of the joints are shown in Figure 3. For the successful trials, the shoulder contributed the most movement for 13 trials, the elbow the most for 4 trials, and the wrist the most for 2 trials.

**Figure 3.**
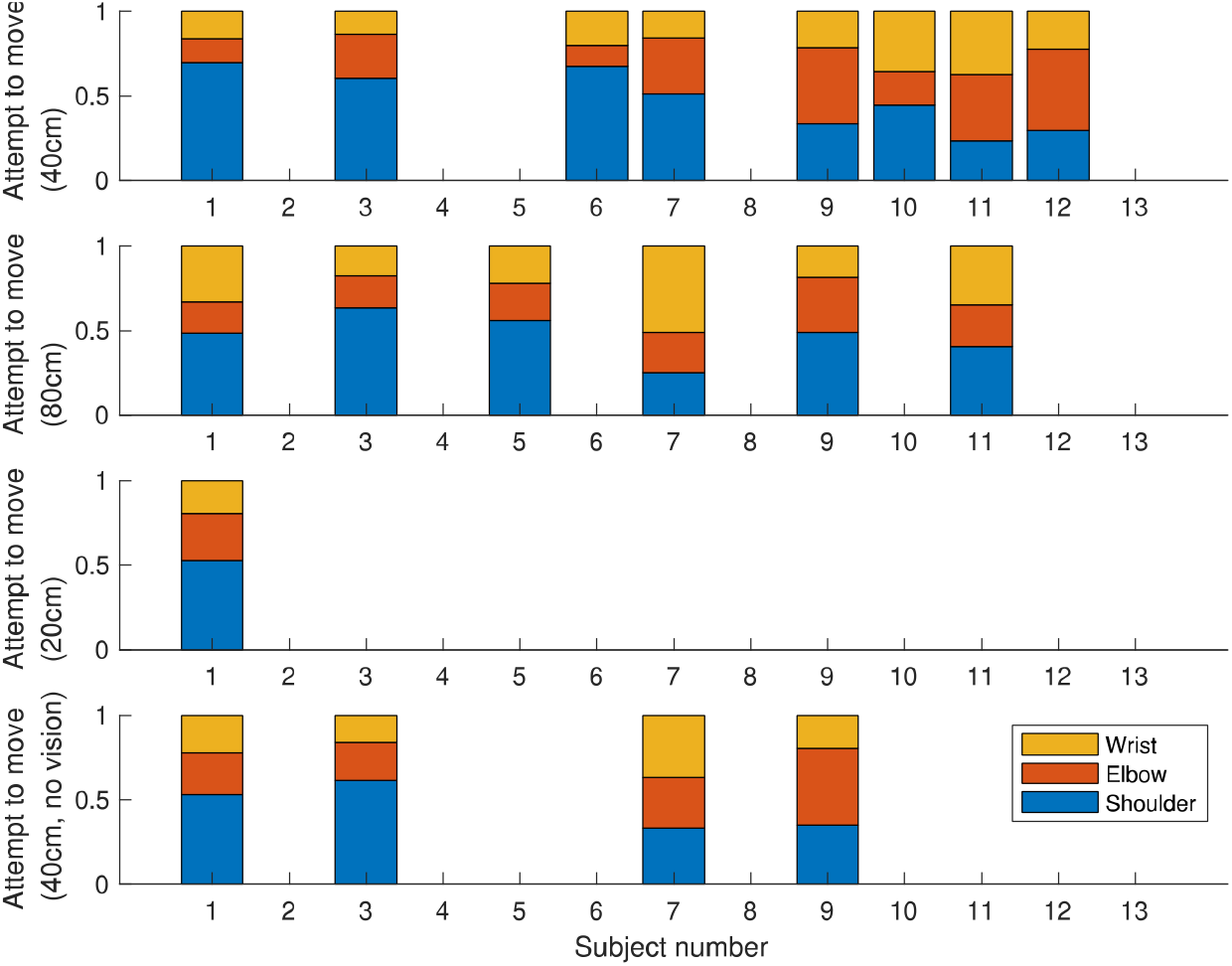
Relative contributions of the wrist, elbow and shoulder to left-right (y) movements of the pendulum

## Discussion

Hand-held pendulums can seemingly oscillate on their own, without perceived conscious control. This phenomenon, known as the Chevreul pendulum illusion, is likely a result of subtle muscle movements caused by thinking of the generated movement. In this study, we examined how the hand-held pendulum, at three different resonant frequencies, results in significant involuntary oscillations which drive the pendulum movement. We demonstrated that the Chevreul pendulum illusion is produced by oscillating the fingers holding the pendulum at a frequency close to the resonant frequency of the pendulum. At an appropriate frequency, very small driving movements of the arm were sufficient to produce relatively large pendulum motion, when the pendulum was sufficiently long (40cm or 80cm) but not for a 20cm pendulum. Subjects who tended to move their fingers more even without holding a pendulum were more likely to be successful in producing the illusion. Different subjects used different strategies, although among the successful subjects, the shoulder contributed the largest amount to the oscillations of the pendulum.

Subjects that move more (also without a pendulum) were more successful - perhaps, in agreement with Chevreul’s proposal, as they move more, the pendulum is likely to start moving, and then they reinforce these movements. In a letter to Ampere, Chevreul argued that tendency of movement in a specific direction is caused by the attention on the pendulum, that is reinforced by sight of the holding hand which extends the movement (oscillations) further. In many other cases, like bowling or billiards, while following a movement with our eyes, we tend to move our bodies in the direction we want the moving body to follow, as if directing the movement towards the goal (Spitz & Marcuard, 2001). The observation that subjects with higher movement show greater likelihood of pendulum oscillation agrees with the bi-directionality relationship between thought and action (i.e. thought of movement resulting in actual movement for subjects), consistent with their expectations (Shin et al., 2010). According to Koch, Keller & Prinz (2004), the anticipation of response effects likely serves as a mental cue to activate corresponding movements.

The resonant frequency of these pendulums allowed the very small movements of the hand to be magnified in the movements of the pendulum, where they could be cumulatively built upon until a regular swinging motion ensued. Why were many participants successful at 40cm and 80cm and not 20cm? The resonant frequency for 20cm is relatively high (2 Hz) but still lower than tremor, similar to frequency of typical movements (whereas the others are relatively slow). It is unclear why it seems difficult to unconsciously produce movements at this frequency. A possible explanation is that the 40cm and 80cm pendulums have resonant frequencies (1.05 Hz and 0.7 Hz respectively) that are close to the natural frequency of the arm, which is approximately 1 Hz (Wagenaar & van Emmerik, 2000), whereas the 20cm pendulum has a resonant frequency (2 Hz) which is far from the resonant frequency of the arm. We note that subjects were more successful with the 40cm pendulum, which is closer to the resonant frequency of the arm.

Some researchers (Karlin, Hill, & Messer, 2007; Olson, Jeyanesan, & Raz, 2017) posit that ideomotor phenomena have large interpersonal variability, wherein the hand-held pendulum oscillations are caused due to different decision strategies. Karlin et al. (2007) showed lower hypnotic susceptibility in subjects that could not produce the Chevreul’s Pendulum illusion, while Olson et al. (2017) showed one of the personality measures, transliminality (sensitivity to subtle stimuli) predicted pendulum performance.

Studying a group of Ouija enthusiasts in a field experiment, Anderson et al. (2019) reported that a combination of retrospective inference and an inhibition of predictive processes caused a reduced sense of agency (subjective experience of not moving the planchette themselves) in Ouija board believers. Herbort & Butz (2012) proposed a computational model of ideomotor theory arguing that action-effect associations are boosted when acting in an *intention-based* action mode. Our findings, that success with pendulum oscillations was twice as high compared to when no visual feedback was allowed, are in line with ideomotor theories that action intentions are essentially perceptual. Mentalizing the movement results in higher success rate when salient visual cues are available.

Rather than a response to a sensory stimulus, ideomotor actions are unconsciously initiated and the results here also demonstrate that priming or thinking of motion can induce muscle micromovements (not visible to the naked eye) that end up moving the pendulum. In light of contemporary ideomotor theory (Shin et al., 2010), knowledge of stimulus-response compatibility is a critical component in pendulum illusion. An action is automatically associated with its effect and that anticipation of the effect facilitates action in a bidirectional manner. Ideomotor actions underlie similar phenomena where items reportedly move of their own accord, like dowsing, facilitated communication or automatic writing, as well as in planchette motions/Ouija phenomenon. In these cases, our intentional stance and sense of agency is likely diminished (Wegner, Fuller, & Sparrow, 2003).

Following on from a scientific tradition of explaining the mechanisms behind seemingly supernatural phenomena (Pfungst, 1911), in this study we demonstrated in detail for the first time the mechanism behind the Chevreul pendulum illusion. This leads to many further questions related to the unconscious production of movement, for example, whether there is a limit on frequencies of movement that can be produced without conscious awareness, and whether the movements produced are a result of prior physics knowledge or simply a result of trial-and-error. Understanding these phenomena may help explain how other types of unconsciously generated movements are produced.

## Author Contributions

JF and BA developed the study concept. Data collection were performed by DC. JF performed the data analysis. DC drafted the manuscript, and JF and BA provided critical revisions. All authors approved the final version of the manuscript for submission.

## Acknowledgements

We thank Alison Kabo for help in data collection.

